# TRAUMA AND CUE-ASSOCIATED WORKING MEMORY DEFICITS IN A RAT MODEL OF POSTTRAUMATIC STRESS DISORDER

**DOI:** 10.1101/2022.05.20.492890

**Authors:** C. E. McGonigle, C. C. Lapish, M. L. Logrip

**Author notes:** Corresponding Author Marian L. Logrip, PhD. Department of Psychology Indiana University-Purdue University Indianapolis 402 N Blackford St. Indianapolis, IN 46202.

## Abstract

Posttraumatic stress disorder (PTSD) is associated with a variety of neural and behavioral alterations in response to trauma exposure, including working memory impairments. Rodent models of PTSD have not fully investigated chronic or reactive working memory deficits, despite clinical relevance. The present study utilizes footshock trauma to induce a posttraumatic stress state in rats and evaluates the effect of trauma and trauma-paired odor cues on working memory performance in the odor span task. Results demonstrate the emergence of chronic deficits in working memory among traumatized animals by three weeks post-trauma. The presentation of a trauma-paired odor cue was associated with further decrement in working memory performance. Further, anxiety-like behaviors indicative of PTSD can predict the degree of working memory impairment in response to the trauma-paired odor cue. This study enhances validation of an existing rodent model of PTSD through replication of the clinical observations of working memory deficits associated with PTSD. This will facilitate future work to probe underlying mechanistic dysregulation of working memory following trauma exposure and for future development of novel treatment strategies.

## INTRODUCTION

Posttraumatic stress disorder (PTSD) is a psychological condition characterized by a pervasive maladapted stress state, elicited by a significant trauma. PTSD is defined by four primary symptom clusters: intrusive thoughts and memories, avoidance, negative alterations in mood and cognition, and hyperarousal and reactivity (American Psychiatric Association 2013). Intrusions and changes in reactivity are linked with adverse reaction to internal or external trauma reminders and demand substantial attentional and cognitive resources (Cox and Olatunji 2017). Working memory also is dependent upon these attentional and cognitive resources and is commonly impaired in PTSD patients (Honzel et al. 2014). This relationship has brought about potential interventions in clinical applications where working memory training has been demonstrated to improve symptoms of posttraumatic stress disorder, suggesting an intersection between working memory and PTSD mechanisms (Larsen et al. 2019; McDermott et al. 2016; Saunders et al. 2015). Despite these successes in clinical applications, working memory has not been probed in animal models of PTSD, which will be necessary to provide a mechanistic understanding of the relationship.

### Posttraumatic Stress Disorder

PTSD affects approximately 8% of the population, although this only represents a small percentage of the population that has experienced trauma (Kilpatrick et al. 2013). Little is known about the mechanisms underlying susceptibility to PTSD, but it is clear that not everyone exposed to significant trauma will develop PTSD. Susceptibility also is accompanied by significant sex differences. Despite being less likely to experience trauma in their lifetime, women are nearly twice as likely as men to develop PTSD (Kessler et al. 1995, 2005).

Animal models have traditionally evaluated all traumatized animals as a single, uniform group, limiting the nuanced exploration of susceptibility factors (Richter-Levin et al. 2019). A series of approaches have attempted to mimic clinical diagnostic procedures by employing a battery of behavioral tests selected to approximate symptom clusters to determine whether an animal is presenting with a PTSD-like state (Cohen et al. 2003; Richter-Levin et al. 2019). Multivariate approaches such as this accommodate individual differences in symptom presentation and may better capture sex differences associated with symptom profiles (Ardi et al. 2016; Murphy et al. 2019). Additionally, animal models of PTSD utilize varied trauma procedures, which may further variability in symptom presentation.

Fear conditioning has been a dominant tool for modeling PTSD and generates a phenotype of lasting reactivity toward cues and context associated with the stressor, most often footshock (Bienvenu et al. 2021). Footshock trauma has strong face and predictive validity for PTSD, resulting in behavioral alterations akin to the primary symptom clusters of clinical PTSD and responding to pharmaceutical interventions for PTSD (e.g. Paroxetine (Paxil) and Fluoxetine (Prozac)) (Bali and Jaggi 2015). Footshock trauma meets another important criterion of a valid PTSD model in that the behavioral responses persist and/or progress with the passage of time (Mikics et al. 2008; Yehuda and Antelman 1993). Footshock generally is not administered at painful levels, but its success as a trauma model relies upon the ability for surprising, unpleasant stimuli to induce long-lasting behavioral alterations (van Dijken et al. 1992; Wellman et al. 2014).

Methods of modeling PTSD-like trauma have largely focused on replicating physiological and reactivity phenotypes of PTSD (Cohen et al. 2012; Liberzon et al. 1997). However, additional assessment is needed of the trauma memories at the root of these physiological and reactivity profiles. PTSD has a multifaceted relationship with memory, defined by overactive and inappropriate memories of the trauma, but also commonly resulting in generalized memory impairments, including working memory (Nejati et al. 2018).

### Working Memory

Working memory is a limited-capacity mechanism for temporarily retaining and manipulating information for use in goal-directed tasks (Barch et al. 2009). Stress and trauma create chronic energetic and attentional demands that reduce available resources for normal functioning. This diversion of attentional resources may provide a mechanism to understand working memory deficits in PTSD (Balderston et al. 2017; Block and Liberzon 2016; Peters et al. 2017). PTSD has been associated with elevated levels of attentional bias to aversive stimuli and it is theorized that the persistence of trauma-related memories serves as a low-level distractor that is omnipresent for the affected individual (Woodward et al. 2017). Clinical studies have demonstrated working memory deficits following acute stress, as well as in persistent states of stress such as PTSD (e.g. Moran 2016; Scott et al. 2015; Shields et al. 2016). Rodent models have replicated the effects of acute stress on working memory performance but have not been utilized to study the effect of long-lasting stress responses that characterize PTSD on working memory (Davies et al. 2013). Span tasks, such as the odor span task, are the primary method for testing working memory capacity in rodents and therefore optimal to determine the effects of stress on this behavior (Davies et al. 2013; Dudchenko 2004; Dudchenko et al. 2013).

The present study utilized the odor span task to assess working memory performance in a rodent model of PTSD. Rats underwent footshock trauma in the presence of a trauma-paired odor (TPO), and subsequent working memory performance was evaluated in the presence that TPO. Two hypotheses were tested: that working memory performance would be impaired acutely and/or chronically; and that working memory performance would deteriorate when presented with the TPO in the odor span task.

## RESULTS

Rats were assessed for the impact of trauma on both anxiety-like and working memory behaviors, as illustrated in Figure 1A. To examine working memory, an odor span task was utilized, in which rats identified novel odors in an incrementing delay non-match-to-sample (DNMS) task (Figure 1B), as described in Materials & methods. Prior to beginning odor span training, animals went through a pre-test elevated zero maze test to evaluate baseline anxiety-like behavior (Fig. 1C). Following two weeks of initial training, DNMS performance was evaluated to assess baseline task performance (Fig. 1D). Animals were counterbalanced into control and trauma groups on these scores. Anxiety-like behavior, as assessed by time spent in the open zones of the zero maze, did not differ by condition [F(1,28) = 0.0050, p = 0.94], but there was a main effect of sex [F(1,28) = 5.33, p=0.024]. Females spent more time exploring the open zones of the maze, which is often observed in locomotor-based tasks. There were no significant differences in DNMS performance between the control and trauma conditions [F(1,28) = 0.27, p = 0.61] or sexes [F(1,28) = 2.07, p = 0.16].

**Figure 1.**
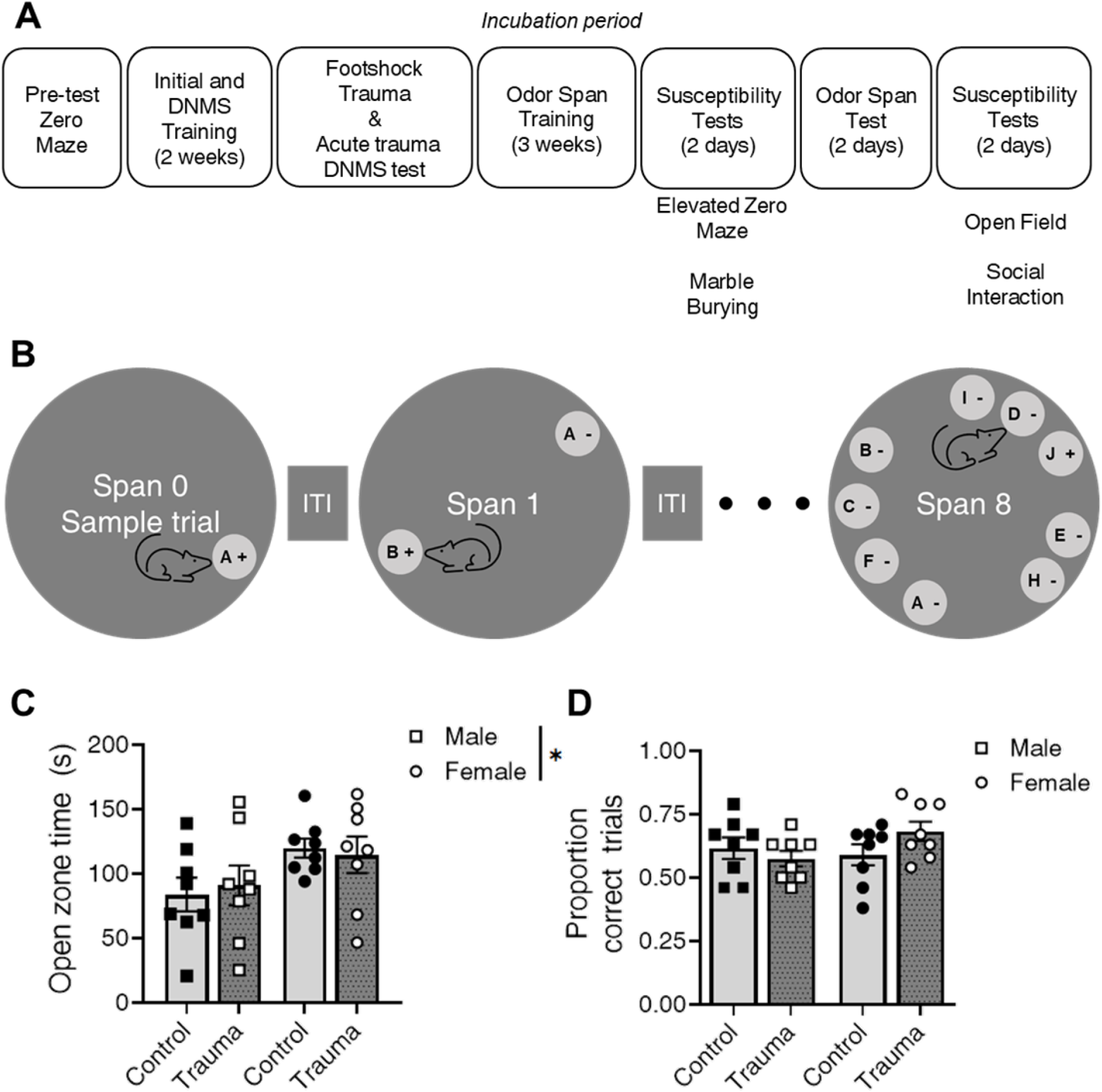
Experimental procedures and counterbalancing measures. A. Experimental timeline. ITI = Intertrial interval. B. Depiction of sample odor span session. C. Group assignments show balanced pre-test elevated zero maze performance, as assessed by open zone time, although females spent greater time in the open zones of the maze than males. *p < 0.05, male vs female. D. Pre-stress DNMS performance did not differ between the groups. DNMS = Delayed non-match-to-sample. N = 8 per group.

Working memory performance in the DNMS training phase was assessed immediately following the footshock trauma session. Data are expressed as percent change in correct responses compared to the previous DNMS session (Fig. 2). There were no group differences between male and female animals [F(1,28) = 0.64, p = 0.43], control and trauma animals [F(1,28) = 0.18, p = 0.68], or interaction between sex and condition [F(1,28) = 0.043, p = 0.84]. No change in DNMS performance was observed following acute stress for any individual group (male control [t(7) = 1.96, p = 0.090]; male trauma [t(7) = 1.32, p = 0.23]; female control [t(7) = 1.11, p = 0.30]; female trauma [t(7) = 1.33,p = 0.23]).

**Figure 2.**
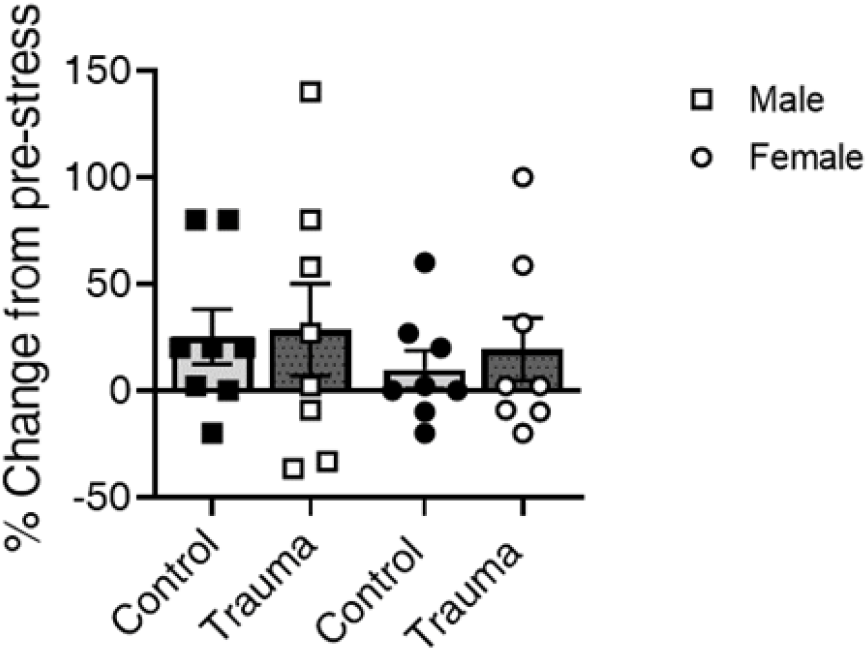
Delayed non-match to sample change following acute stress. Acute stress did not significantly impact DNMS performance, expressed as a change score calculated as percent change from DNMS performance pre-stress. N = 8 per group.

Working memory performance was assessed during the final week of training as an average measure of baseline performance (Fig. 3A). There was a main effect of sex [F(1,28) = 4.87, p = 0.036], demonstrating that female working memory performance was, on average, higher than male performance. There was a main effect of condition [F(1,28) = 4.76, p = 0.038], revealing that animals with a trauma history obtained lower maximum spans, on average, than control animals. There was no interaction between sex and condition [F(1,28) = 0.19, p = 0.66]. Omissions also were summed from the final week of training to examine whether the lowered trauma-history spans could be explained by animals prematurely “quitting” trials through omissions (Fig. 3B). Two-way ANOVA revealed no main effect of condition [F(1,28) = 2.42, p = 0.13], supporting the fact that trauma-history animals did not omit responses significantly more often than control animals. There was no main effect of sex [F(1,28) = 0.097, p = 0.76], suggesting that the sex difference in working memory performance was not driven by an increase in trial omissions by males.

**Figure 3.**
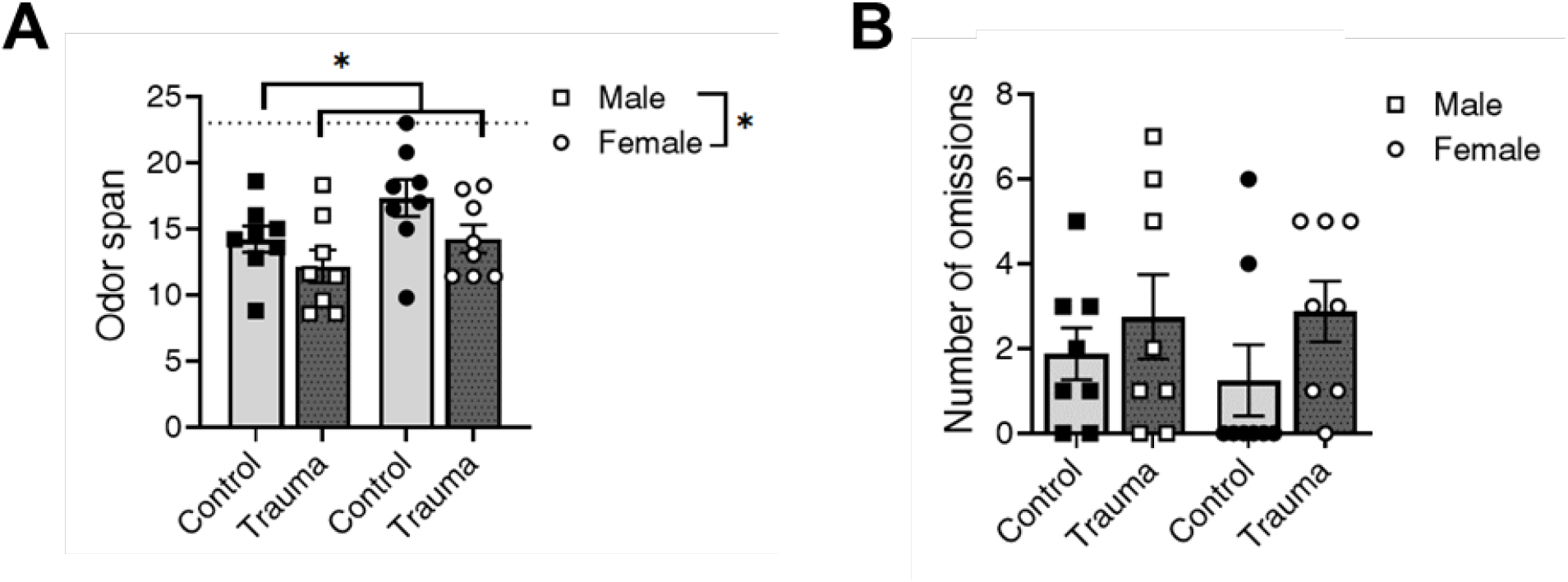
Trauma history impacted training performance in the odor span test. A. Maximum span, calculated as the average maximum odor span across the final five days of training, was significantly lower in males (squares, left) and animals with a trauma history (dark bars, right in each pair). B. Summed trial omissions from the final five days of training did not differ between groups. N=8 per group. * indicates male vs female and control vs trauma (p < 0.05).

All animals received two days of behavioral testing, elevated zero maze and marble burying, discussed below, and then entered the testing phase of the odor span task. In this phase, animals were given a normal testing day to serve as baseline and a TPO testing day. On the TPO testing day, the TPO, coriander, was presented as the target cup at span 4. Maximum span performance is presented for baseline and TPO testing days (Fig. 4A). Percent change in span length on the TPO session was calculated as a metric of how each animal’s working memory performance was affected by the presence of the TPO (Fig. 4B). One-sample t-tests revealed that male control [t(7) = 0.72, p = 0.49] and female trauma [t(7) = 0.69, p = 0.51] animals did not have significantly altered spans on the TPO test session. On the other hand, male trauma [t(7) = 2.82, p = 0.026] and female control [t(7) = 3.30, p = 0.017] animals demonstrated significant decreases in maximum span on the TPO session.

**Figure 4.**
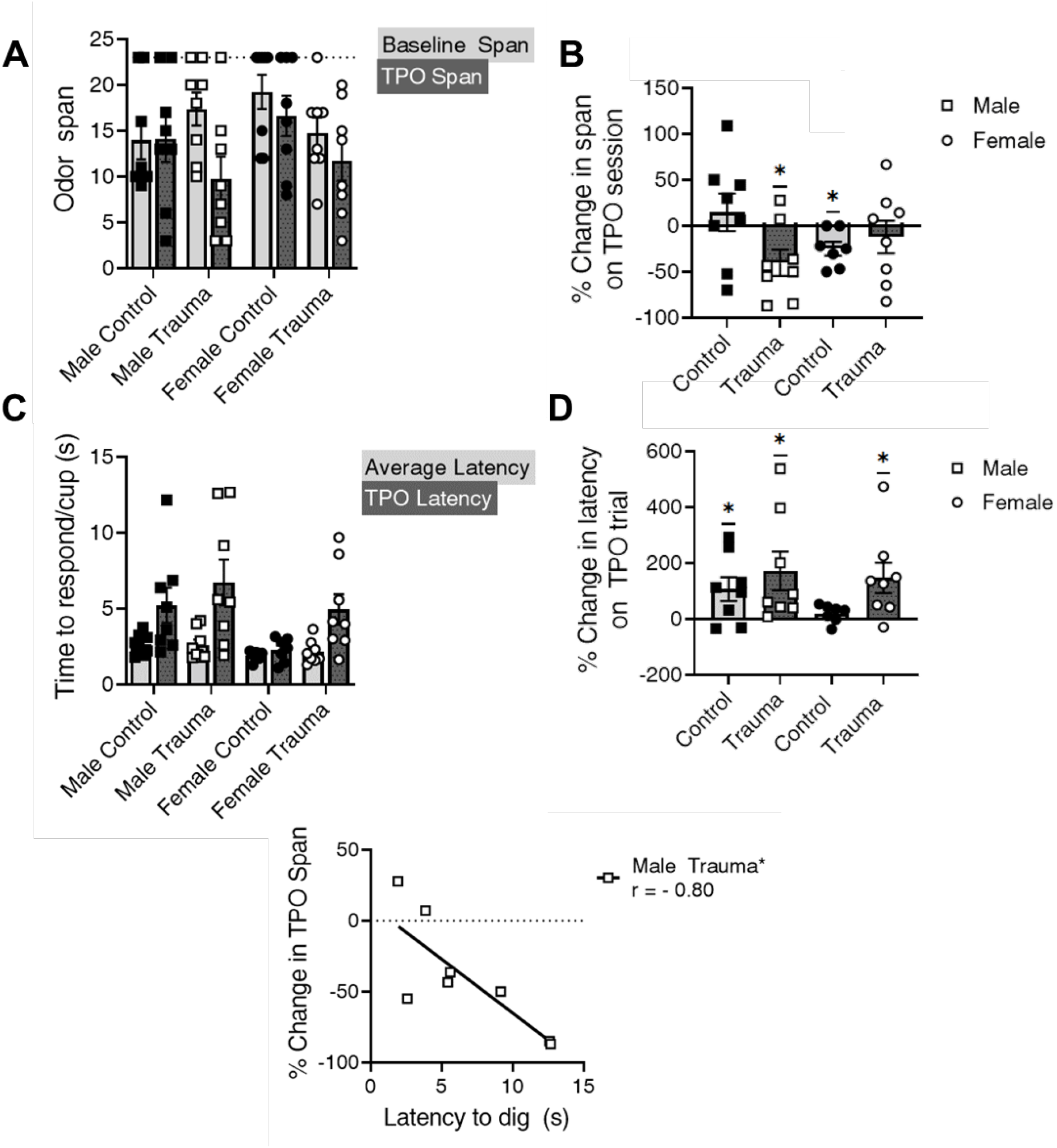
Trauma history impacted performance in the odor span test. A. Data show maximum span obtained on baseline test day (left bars) and trauma-paired odor (TPO) test day (right bars). B. Span change on the trauma-paired odor test day was significantly different from baseline for male trauma and female control animals. * p < 0.05, one-sample t-test compared to no change (0). C. Latency to respond on non-TPO trials, calculated as the average latency per sampled cup on choice trials 1-3, compared to TPO trial 4. D. Latency to dig on the TPO trial increased for all groups except female controls * p < 0.05, one-sample t-test compared to no change (0). E. Latency to dig on the TPO trial correlated with span impairment only for male trauma animals, lines correspond to group correlation slopes * indicates p < 0.05, male trauma. N=8 per group.

Latency to dig is presented for non-TPO trials and the TPO trial, calculated as the total latency to dig divided by the number of cups sampled (Fig. 4C). Percent change in latency to dig was calculated to directly compare the change in latency to respond within each group (Fig. 4D). Female control animals did not significantly increase latency to dig on the TPO trial [t(7) = 1.68, p = 0.14], but male control [t(7) = 2.50, p = 0.041], male trauma [t(7) = 2.50, p = 0.041], and female trauma [t(7) = 2.71, p = 0.030] animals all significantly increased latency to dig on the TPO trial. Deficits in span were hypothesized to result from cue reactivity, which is approximated by latency to dig on the TPO trial. Interestingly, the latency to dig on the TPO trial was only significantly correlated with the decrement in span performance for male trauma animals [Pearson’s r = −0.80, p = 0.0017]. All other groups demonstrated non-significant relationships between latency and span change (male control [r = 0.099, p = 0.82]; female control [r = 0.10, p = 0.83]; female trauma [r = −0.35, p = 0.40]).

Susceptibility tests were run before and after the odor span testing days to protect against the possibility that re-exposure to the trauma-paired cue substantially affected the behavior of the trauma-history animals. Elevated zero maze behavior was measured by time spent in the open zones of the maze (Fig. 5A). Two-way ANOVA revealed no main effect of condition [F(1,28) = 0.11, p = 0.74] or sex [F(1,28) = 0.070, p = 0.79] on open-zone time. General locomotor activity was evaluated via the number of entries into the open zones during the elevated zero maze test (Fig. 5B). There was no main effect of condition [F(1,28) = 0.0049, p = 0.94], but there was a main effect of sex [F(1,28) = 19.13, p = 0.0002], demonstrating that females displayed greater general locomotion during the zero maze test even though their total time in the open zones did not differ from males. Marble burying results are not shown because only 3 trauma-history and 1 control out of the 32 total animals engaged in marble-burying behavior. Open field behavior was scored as the total amount of time that the animal spent in the center of the arena (Fig. 5C). There were no main effects of condition [F(1,28) = 0. 025, p = 0.88] or sex [F(1,28) = 0. 48, p = 0.49] on center time. Total distance traveled, scored with ANY-Maze as number of zone crossings multiplied by the size of each zone in the maze (15 cm), was analyzed for locomotor behavior (Fig. 5D). There was no main effect of condition [F(1,28) = 0.078, p = 0.78], but a trend toward a main effect of sex [F(1,28) = 3.99, p = 0.056], again supporting that females tend to have higher levels of general locomotion than males during anxiety-like behavior tests. Behavior in the social interaction test was evaluated by the distribution of time spent engaged in nonsocial, prosocial, and antisocial encounters (Fig. 5E). A priori hypotheses were that a PTSD-like phenotype would be associated with less total social interaction and greater engagement in antisocial behavior. Two-way ANOVAs were performed on each of these metrics. Total social interaction did not differ by condition [F(1,28) = 0.21, p = 0.65], but there was a main effect of sex [F(1,28) = 8.74, p = 0.0063], demonstrating that females spent less total time engaged in social interaction. There were no main effects of condition [F(1,28) = 0.045, p = 0.83] or sex [F(1,28) = 2.43, p = 0.13] in time spent engaged in antisocial behavior. Data were converted to z-scores within each group (male control, male trauma, female control, and female trauma) to allow for combination of male and female data by condition, while accounting for baseline differences between the sexes. When normalized within condition, the change in span on TPO trials is significantly correlated with antisocial behavior for trauma-history animals only (Fig. 5F; Pearson’s r = 0.64, p = 0.007). Control animals did not display a significant relationship between antisocial interaction and change in span (r = −0.38, p = 0.16).

**Figure 5.**
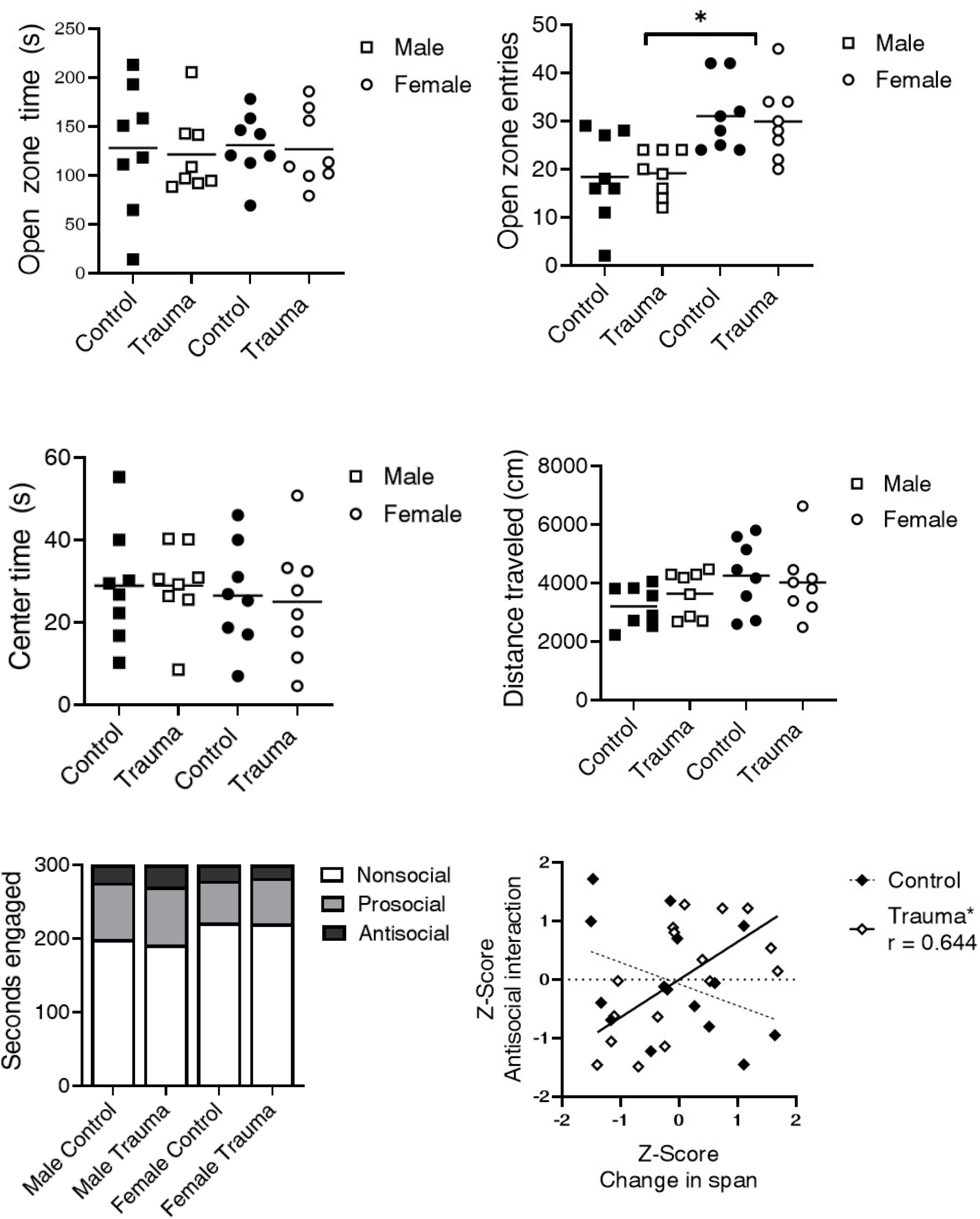
Behavioral profiling to assess PTSD-like phenotype. A. Elevated zero maze open zone time did not differ between groups. B. Open zone entries in zero maze were more numerous for females, indicating greater locomotor activity. *p < 0.05, male vs. female. C. Open field time in the center zone did not differ between groups. D. Total distance traveled in the open field test did not differ between groups. E. Time spent engaged in nonsocial, prosocial, and antisocial behaviors in the social interaction test differed by sex. F. Correlation between normalized change in span and antisocial interaction was significant only for animals with trauma-history, lines indicate group correlation slopes. *p < 0.05, control vs. trauma. N=8 per group.

Multiple regression analysis using the PTSD-like behavioral profile significantly predicts the degree to which an animal’s span was altered by presentation of the trauma-paired cue (span change) for trauma-history animals. For this analysis, data from each predictor variable (training span, TPO latency, elevated zero maze open zone time, open field center time, and antisocial interaction time) and the outcome variable (percent change in span) were converted to z-scores to standardize data scaling across groups and collapsed across sex since few effects of sex were present in the data. This model was significant for trauma-history animals [Table 2; F(5,10) = 18.12, p < 0.001, R^2^ = 0.90, adjusted R^2^ = 0.85, all variables contributed significantly to the prediction, p < 0.05], but did not differ from the null for control animals [Table 1; F(5,9) = 1.023, p = 0.46, R^2^ = 0.36; no variables contributed significantly to the prediction, p > 0.05]. Bootstrapping results confirmed the validity of the point-estimates for the regression coefficients (Table 2). Repeated 5-fold cross validation was performed on the trauma-history model, resulting in an average prediction error (root mean squared error) of 0.48. Covariance matrices are presented in Figure 6.

**Figure 6.**
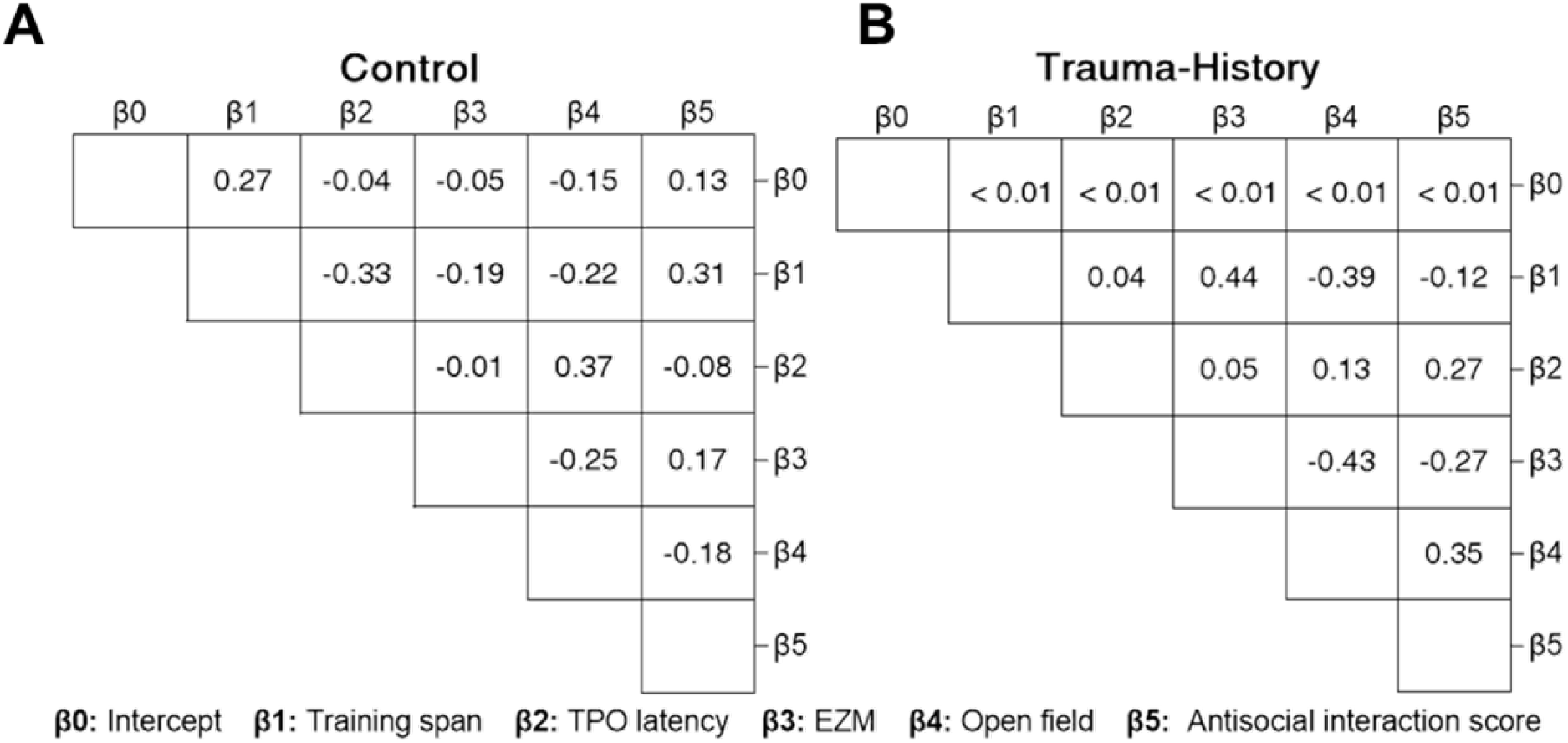
Multiple linear regression covariance matrices. No regression parameters significantly covaried for control animals (A) or trauma-history animals (B).

**Table 1.**
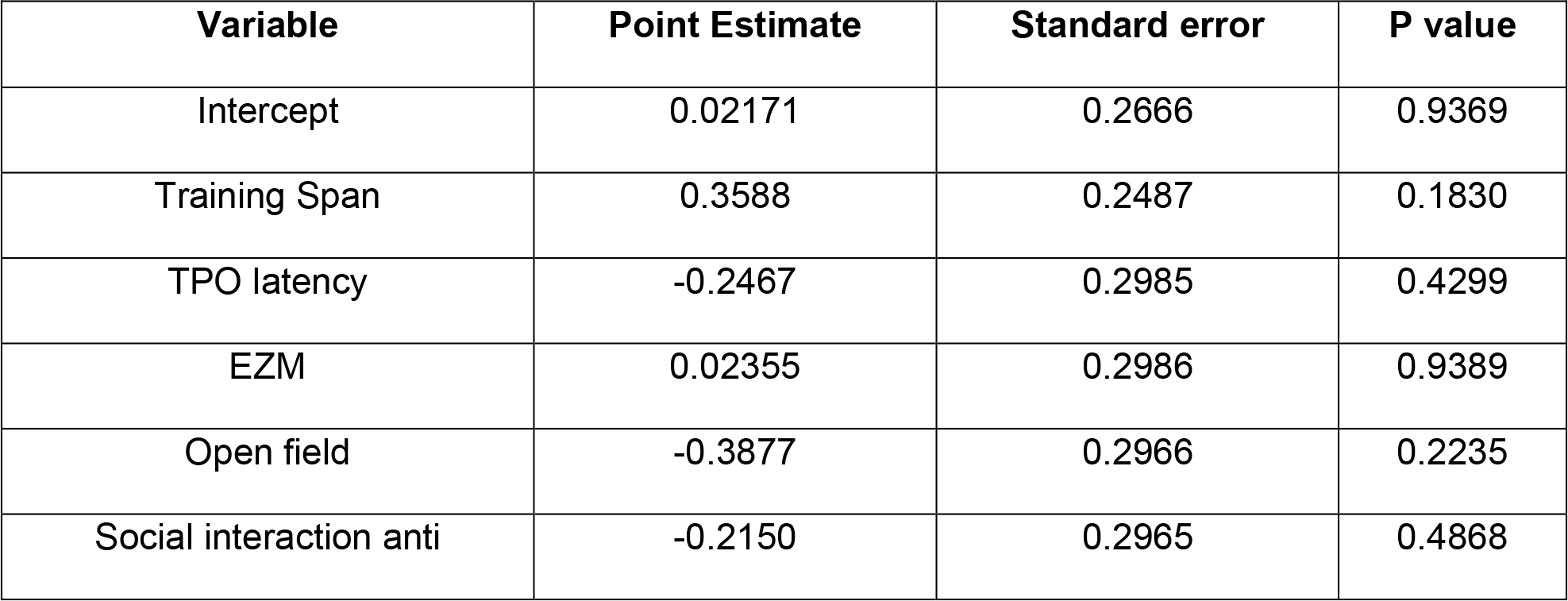
Multiple regression for control animals.

**Table 2.** Multiple regression for trauma animals. Bootstrap CI indicates 95% confidence interval, calculated using 2000 bootstrap replicates. * indicates p < 0.05; ** indicates p < 0.01; *** indicates p < 0.001.

| **Variable** | **Bootstrap CI** | **Point Estimate** | **Standard error** | **P value** |
| --- | --- | --- | --- | --- |
| Intercept | (-0.25 - 0.23) | 4.764e-007 | 0.09327 | >0.9999 |
| Training Span | (0.28 - 0.93) | 0.5549 | 0.1153 | 0.0007*** |
| TPO latency | (-0.69 - 0.14) | -0.4124 | 0.1050 | 0.0028** |
| EZM | (0.27 - 0.87) | 0.5439 | 0.1195 | 0.0011** |
| Open field | (-0.63 - 0.01) | -0.3133 | 0.1194 | 0.0255* |
| Antisocial interaction | (0.04 - 0.62) | 0.3929 | 0.1116 | 0.0055** |

## DISCUSSION

The present study is among the first to examine working memory in an animal model of PTSD. A footshock trauma procedure and the odor span task were used to evaluate the effect of trauma history and trauma-paired cue reactivity on working memory performance. The data demonstrated that while footshock trauma did not acutely affect working memory performance in DNMS (Figure 2), trauma history was associated with persistent deficits in working memory performance (Figure 3), and exposure to a trauma-paired cue further impaired working memory (Figure 4). Additionally, multivariate analysis of PTSD phenotypic behaviors significantly predicts the degree to which an individual’s working memory is impaired by the presentation of trauma-associated cues further validating the change in span as an index that models certain features of PTSD. Collectively, these data move the PTSD field toward an animal modeling approach that will allow for investigation of the mechanisms and implications for working memory deficits in PTSD.

### Lack of acute effects of footshock trauma

Animals completed footshock sessions one at a time, after which they were transported immediately to the behavioral testing room, where they performed 8 DNMS trials to assess the acute impact of trauma. There was no effect of acute trauma on DNMS performance, which appears to contrast with evidence that acute stress impairs working memory capacity in the odor span task (Davies et al. 2013). It is possible that the lack of acute stress effects is driven by differences in mechanisms (or resources required) for working memory capacity versus two-sample discrimination. It may not be surprising to find such conflicting results, as acute stress has been found to both impair and enhance working memory performance in different working memory tasks, and whether performance is impaired or enhanced may be partially dependent on the sex of subject (Khayyer et al. 2021; Schoofs et al. 2013). Further, in the present study, animals were still in a training phase and the lack of established stable responding may have obscured the true effect of acute footshock trauma on working memory performance.

### Past trauma triggered chronic working memory deficits

Following footshock trauma, animals were given three additional weeks of odor span training, which also served as a stress-incubation period to allow for the emergence of persistent stress phenotypes (e.g. Harvey et al. 2003). Maximum odor span was averaged across the final week of training, and trauma history resulted in lower overall working memory performance at three weeks post-trauma compared to the control group. Both sexes with trauma history had lowered maximum spans compared to controls, although females had higher spans regardless of condition. Clinical research indicates that females may have slightly better working memory performance than males, but these results may be impacted by factors such as general motivation and attention or olfactory sensitivity (Sanchez-Andrade et al. 2005; Voyer et al. 2021). It should be noted that span performance in this study was high compared to previous odor span studies (Davies et al. 2013; De Falco et al. 2019; Dudchenko et al. 2000). The most apparent methodological differences between past odor span work and the current study were the present study’s use of Wistar rats and inclusion of both sexes. Previous studies used Long Evans and Sprague Dawley strains, and primarily focused on males. It is possible that working memory, stress response, olfactory sensitivity, or general cognitive performance may be confounded by strain and sex (Andrews 1996; Martis et al. 2018). Probe trial performance indicated that animals were not marking cups or selecting based on the smell of the buried food reward, but that does not rule out possible engagement in strategies like chunking in a manner that promoted attainment of higher spans (Guida et al. 2012).

### Trauma-paired cue reduced span performance for trauma-exposed rats

Following the three weeks of OST training and stress incubation, animals performed a baseline odor span test, which was identical to training days. On the next day, they were given a TPO test, in which the TPO was presented at span 4. Male trauma-history animals had significantly lower span performance when the TPO was present. Female trauma-history animals did not demonstrate a significant deficit in span in the presence of the TPO, although this may have been driven by greater variability among trauma-history females. The apparent working memory deficit in female control animals in the presence of the TPO is likely driven by the fact that many of these females were regularly attaining the maximum span possible in the test, and so a small variation from this ceiling was significant. Average latency to dig was significantly higher on the TPO trial in all groups but female control, likely indicating deliberation or hesitancy to interact with the TPO. Male trauma history animals are the only ones for whom latency to dig on the TPO trial was significantly correlated with the deficit in span performance in the presence of the TPO. This correlation was not driven by animals who were unable to continue the span following the TPO trial. Rather, animals who successfully completed the TPO trial subsequently achieved a lower span than at baseline.

The effect of the TPO on working memory demonstrates cue reactivity in the current model, which is an important aspect of the PTSD-like phenotype. Cue reactivity is associated with two of the primary PTSD symptom clusters: intrusions and hyperarousal/reactivity. The disruption of task performance in the presence of the TPO may indicate that the animals were distracted by its presence. Attention plays a central role in the working memory mechanism, and distractors generally impair working memory performance (Cowan 2008). A chronic response to trauma may involve allocation of attentional resources to trauma-associated information (Peters et al. 2017). Therefore, the working memory deficits following exposure to trauma cues may be related to impairments in the ability to maintain attention on the task when the trauma-paired odor is present. Indeed, targeting attention maintenance and working memory deficits in individuals diagnosed with PTSD reduced their symptoms (Badura-Brack et al. 2015; McDermott et al. 2016).

Odor span is considered to capture working memory capacity, which is conceptually distinct from the virtually limitless recognition memory (Turchi and Sartor 2000; Dudchenko 2000). Span tasks require than an animal recall the order of stimulus presentation, which assesses working memory rather than recognition memory. It has been shown that working memory capacity frequently falls into the range of 7 +/- 2 items in humans and animals, although smaller limits have been proposed (Cowan 2001; Hahn et al. 2021; Miller 1956). Within the odor span task, performance remains high when more than 70 odors are to be recognized if the maintenance of presentation order is not required (April et al. 2013). Unique patterns of neural activity have been observed to support recognition of “novel” versus “familiar” odors, suggesting that the odor span task may not require the memory of each individual odor (De Falco et al. 2019). These may suggest that the odor span task captures elements of cognitive performance beyond working memory. Recognition and working memories are likely both dependent upon attentional allocation and may be impacted by trauma history, as attentional deficit and bias have been associated with impairment in both forms of memory (Fosco et al. 2020; Spataro et al. 2022). Thus, future work must expand on the current findings, particularly by examining the mechanisms underlying attentional allocation toward trauma cues and manners through which the resulting attentional bias may be corrected in animal models.

### Multivariate prediction of cue reactivity

Animals were tested on elevated zero maze, marble burying, open field, and social interaction to evaluate PTSD-like phenotypes. PTSD is a multifactorial disorder characterized by clusters of symptoms; therefore, a multiple linear regression model was built to determine whether this array of behavioral indices could predict how trauma cues impact working memory performance (change in span). Significant predictors included working memory span performance during training, latency to dig in the TPO trial, elevated zero maze open zone time, open field center time, and antisocial behavior. For animals with a trauma history, this combination of behaviors held significant predictive value for the degree to which trauma presentation would impact working memory performance. These results suggest that trauma-paired cue reactivity is associated with working memory deficits and relates to a wider profile of PTSD-like symptoms.

Previous studies have employed strategies with fewer measures (generally two tests) of anxiety-like behavior to determine extent of the PTSD-like phenotype in animals. Clinical diagnosis of PTSD requires symptoms across four primary clusters, and male and female symptoms may differ, so the utilization of multiple tests is important to capture individual variation (Carmassi et al. 2014; Carragher et al. 2016; Gruene et al. 2015). Future work will be needed to validate the use of a multiple linear or logistic regression approach to detect susceptible and resilient phenotypes in animal models of PTSD. Additionally, the current study relied upon multiple behavioral endpoints, which are inherently variable. Larger sample sizes would provide additional power for detecting significant effects, particularly with respect to sex differences and susceptible phenotypes. To this end, future studies would benefit from utilization of higher throughput model, such as an automated odor span task (Galizio et al. 2020). Additionally, because male and female rodents exhibit differing anxiety-like behavioral profiles, future work should incorporate anxiety-like tests designed to capture more female-typic behaviors (Gruene et al. 2015; Meyerson et al. 2006).

## Conclusion

PTSD models lack standardization, resulting in variations in models associated with differing behavioral outcomes (Bienvenu et al. 2021; Knox et al. 2012). The current study presents a model for PTSD-like impairment of working memory performance, with different impacts in the absence versus presence of a trauma cue. Additionally, the regression model produced a quantitative assessment of the impact of the trauma-paired cue on span performance. The sample size used in the present study was quite small for use in multiple linear regression. Bootstrapping and k-fold cross validation methods were used to provide assurance of model robustness, but the risk remains that these models built on small samples will not generalize well. This method should be validated using a larger sample size to provide further confidence about the ability for a holistic PTSD-like phenotype to predict reactive impairments in working memory. Alternate PTSD models must validate working memory phenotypes to identify molecular underpinnings of these persistent effects of trauma, as many of the current models have not tested this behavioral phenotype. Identification of these molecular factors may yield treatments with increased efficacy in individuals suffering PTSD-related working memory deficits.

## MATERIALS AND METHODS

### Subjects

Three separate cohorts of adult male and female Wistar rats (initial weights M: 253-306 g; F: 146-206 g Charles River, NC) were obtained at 8 weeks of age. Upon arrival, the rats were pair-housed with lights on a 12-hr light/dark cycle (lights on at 0600) with ad libitum access to food and water. Subsequently, subjects were individually housed, mildly food restricted to maintain 90% of free feeding weight, and handled 5 minutes each day for 5 days prior to the first experimental task. All experimental procedures were conducted during the light phase of the light cycle. All procedures were approved by the IUPUI School of Science Animal Care and Use Committee.

### Experimental Overview

The experiment consisted of the odor span task, footshock trauma, and susceptibility tests (Fig. 1A). All behavioral apparati were cleaned with water between same-sex subjects and with 70% ethanol between sexes to minimize odor distractions. Prior to each anxiety-related behavioral test, animals were given an acclimation period in the antechamber of the testing room. White noise (approximately 54 dB) was played during all behavioral tests and tests were run under normal house lights unless otherwise specified. Behavioral tests were recorded via an overhead camera and scored by an observer blind to experimental condition.

### Odor Span Task

Methods for the Odor Span Test followed Davies et al. (2013). The apparatus featured a grey textured plastic platform (91.5 cm round, 30.5 cm border, 79 cm above the ground). Odors (0.5 g of dried spice; allspice, anise seed, basil, caraway, celery seed, cinnamon, cloves [0.1 g], cocoa, coffee, cumin, dill, fennel seed, garlic, ginger, lemon, marjoram, nutmeg, onion powder, orange, oregano, paprika, rosemary, sage, and thyme) were mixed in Premium Play Sand (100 g; Quikrete Cement Products) in 3.25-oz plastic cups (5 cm tall; 7.6 cm wide). Twenty-four Velcro attachment points were equally spaced along the perimeter of the platform and cups were randomly attached to one of these points in each trial to prevent the use of spatial strategies. Animals were trained five days each week in either a morning group or an afternoon group, with sexes and conditions represented equally across groups.

### Initial Training

Initial training involved rats learning to dig for a cereal reward (1/4 of a Kellogg’s Froot Loops®) in unscented sand (3.25-oz cups containing 100 g of sand). For each trial in this phase, a single cup of unscented sand was placed in a random position on the platform. After the subject retrieved the reward or following a maximum response time of 2 minutes, the subject was removed from the platform and placed in a separate intertrial box for a 40-second period. The experimenter then moved the cup to a new location before the next trial. The reward was first placed on the surface of the sand, then buried progressively further in the sand. Animals were trained until they would consistently dig for and retrieve a fully buried reward from the unscented sand. Initial training took approximately 3-4 days.

### Delayed Non-Match-to-Sample

Rats were next trained on a DNMS task. In this phase, rats were first presented with a sample trial where they were presented with a cup of scented sand randomly placed on the platform. After retrieving the reward, rats were moved to the intertrial interval box for a 40-s delay period, during which time the experimenter moved the sample cup to a new location and randomly placed a second bowl containing a different odor and a reward on the platform. The animals were then returned to the platform for their choice trial and allowed to freely sample (sniff) the cups. A choice was determined by the animal contacting the sand with their nose or paws. An error was scored if the rats selected the sample odor, rather than the novel odor. The animals were given 8 DNMS choice trials each day until they selected the novel odor on 6 out of 8 choice trials for three consecutive days. DNMS training took approximately 1.5 weeks.

### Odor Span Task

Following DNMS, animals were trained on the odor span task (Fig. 1B). These trials were conducted identically to the DNMS task, except cups containing novel odors were added following each correct choice until the rat made an incorrect selection or omitted a response, resulting in an increasing number of odors present in the arena. Span for a given trial is defined as the number of choice trials completed successfully, which is equivalent to the total number of cups present minus one. Each rat performed a maximum of three consecutive spans per day, with no trials beginning more than 15 minutes into the session. Maximum span is reported for each day. Animals that reached a stable level of high performance (span > 17 for two consecutive days) were reduced to training three times per week to reduce the likelihood of overtraining.

Following three weeks of OST training, the effect of trauma cue presentation on working memory performance was evaluated. Animals were given a baseline testing day, which was identical to the OST training days except that following an incorrect choice, they returned to their homecage for approximately 30 minutes between spans while other rats were tested. The following day, on span 4 (5 cups present on the table), the TPO (coriander) was presented for the first time as a target odor in the OST task. Latency to interaction with the coriander as well as maximum span achieved while coriander was on the platform were recorded and evaluated.

Several probe trials were conducted throughout training to confirm that subjects were using cup odor to solve the task and to ensure coriander performance could not be attributed to novelty. The first was a non-baited trial to confirm animals were not pursuing the smell of the food reward. At span 4 (five cups on the table), a target cup was presented, which did not contain a food reward. Upon contact with the correct cup, a food reward was dropped onto the surface of the sand for the animals to consume. The second probe trial was a cup-change trial, in which the cups at span 1 (2 cups on the table) were swapped out with clean cups of the same scents. Successful completion of this trial was indicative that the animals were not marking the cups as a strategy to identify previously sampled cups (Davies et al. 2017). Swapping out to clean cups later in the span is not feasible while maintaining a 40-s intertrial interval. Non-baited and cup-changed trials were conducted once per week during the 3-week odor span training period. Lastly, a novel odor probe trial was run with two of the three cohorts to ensure that animals could successfully perform the task when presented with a completely novel odor, confirming that any observed reactions on the TPO test day were not confounded by reactivity to task-novel odors. The novel odor trial was conducted during the final week of odor span training. Performance across all animals on the non-baited probe was 90%, on the cup-change probe was 91%, and on the novel odor probe was 96%.

### Footshock Trauma

Two weeks into the training procedure, animals underwent footshock trauma. They were provided two days of brief acclimation to the footshock chambers prior to the trauma day (<20 min). Animals were counterbalanced into footshock trauma and control groups based on DNMS training performance and initial elevated zero maze performance to maximize chances of equalizing cognitive ability and baseline anxiety-like behavior between the groups. Acclimation and footshock sessions were run in illuminated operant boxes (MedPC), with no additional cues preceding footshock. On the day of the footshock trauma, coriander was placed in the operant boxes, serving as a trauma-paired odor cue, which subsequently was presented in the odor span test to assess the impact of trauma cues on working memory performance. The trauma group had five minutes in the box before footshocks began, followed by 20 uncued shocks at 0.8 mA (1-s duration, VI-40 schedule, resulting in ∼17-minute total footshock session length). Control animals were placed in the boxes with coriander scent present but did not experience any footshock. Similar parameters elicited conditioned fear responses that were sustained three weeks post-stressor, indicating its ability to generate a PTSD-like chronic stress response (Wellman et al. 2014). Animals were shocked one at a time and then immediately brought to the odor span room for that day’s training and assessment of acute stress effects on task performance. Footshock chambers were not used for more than one animal per cohort.

### Elevated Zero Maze

Elevated zero maze performance was evaluated the day following the final odor span training day. This test shows an animal’s response to the conflict between exploration of a novel environment and exposure to an unsafe (open, unwalled) environment, and approximates the avoidance cluster of PTSD-like symptoms (Shepherd et al. 1994; Verbitsky et al. 2020). The maze consists of a circular pathway (105 cm maze diameter, 10 cm track), elevated approximately 50 cm above the floor, with alternating walled and open zones (2 walled and 2 open zones) evenly dividing the total circumference of the track. This circular version of the maze shows fewer sex differences than the plus-shaped version of the maze (Braun et al. 2012; Tucker and McCabe 2017). Testing was conducted in dim lighting conditions (∼0 lux in closed zones, 6 lux on open zones). Rats were placed on the maze facing the entrance of the closed zone and allowed to freely explore for five minutes. Overhead video of the session was captured and scored for time and entries into the open zone, defined by all four paws entering a zone.

### Marble Burying

The marble burying test approximates the PTSD-like symptom cluster of alterations in arousal and reactivity and was conducted on the day following elevated zero maze testing (Kedia and Chattarji 2014; Verbitsky et al. 2020). Marble burying is conducted most successfully in mice, but the tendency to express neophobia through burying behaviors is common among rodent species (Himanshu et al. 2020). Following homecage acclimation to the testing room (normal light, approximately 100 lux), each animal was moved into a standard shoebox cage with 5 cm packed-down bedding and allowed to freely explore for 15 minutes. Then 20 marbles (2 cm diameter) were arranged in a 4 x 5 grid across one half of the testing box (De Brouwer and Wolmarans 2018). The rats were placed on the marble-free side of the testing box and allowed 30 minutes to explore the testing box and interact with marbles, after which they were returned to their home cage and the number of marbles buried was counted. A marble was considered buried if more than ½ was under the bedding.

### Open Field

The open field test provides an index of anxiety-like behavior, as the center of the field is an exposed and unprotected region, and captures PTSD-like avoidance and negative alterations in mood (Gould et al. 2009; Katz et al. 1981; Verbitsky et al. 2020). This test was performed the day following TPO testing in the odor span task. The open field apparatus is a square arena surrounded by high walls (63 x 63 x 50 cm), and testing was performed under moderately dim (40 lux) illumination. The floor of the apparatus is divided into sections to quantify position relative to the walls versus center and movement via line crossings. At the beginning of the test, each rat is placed into the center of the open field arena and allowed to freely explore for 10 minutes. Overhead video of the session was captured and scored for time in the center of the open field and for overall locomotor activity, measured by number of zone crossings multiplied by the size of each zone.

### Social Interaction

This test examines whether rats show deficit in social interactions with an unknown, same-sex conspecific, and approximates negative alteration in mood symptom cluster (File and Seth 2003; Verbitsky et al. 2020). Open field testing was performed the day prior to social interaction testing and served as the acclimation period for the test apparatus. Social interaction was tested under the same lighting conditions as the open field test (approximately 40 lux in the center of the arena). On the test day, rats were paired across condition (one control, one stress) with novel partners. Social interaction most commonly involves pairing animals of identical conditions or pairing with an experimentally naïve animal. However, due to hypothesized individual differences in trauma response, animals were paired across condition and scored individually (Hindley et al. 1985). One rat from each pair was randomly assigned to have a marked tail to allow for individual behavioral scoring during the test. At the beginning of the test, rats were placed in opposite corners of the arena and allowed to interact freely for 5 minutes. All behaviors initiated by the experimental rat are scored according to the following categories: nonsocial behaviors, prosocial and antisocial behaviors towards the other rat.

One day after the social interaction test, animals were euthanized using isoflurane to induce rapid sedation, followed by decapitation.

### Statistical Analysis

Statistical significance was evaluated at p < 0.05 and graphed data are expressed as mean +/- standard error of the mean (SEM) with individual data points overlaid, unless otherwise noted. One-sample t-test was used to evaluate change scores from the null hypothesis of zero change for span and latency data. All groups were independent from one another and therefore no corrections were made for familywise alpha in one-sample t-tests. Two-way ANOVAs were used to compare differences among groups on the basis of sex and condition. Z-score normalization was performed on each group (male control, male trauma, female control, and female trauma) to preserve group differences, and facilitate collapse across sex. Pearson correlation and multiple linear regression were used to assess the significance of bivariate and multivariate linear associations. Bootstrapping (2000 bootstrap replicates) and repeated 5-fold cross-validation techniques (3 replicates) were used to affirm robustness of the multiple regression model. Statistical testing and graphing were performed in GraphPad Prism version 9.3.1 (GraphPad Software, San Diego, California USA) and R (R Core Team, 2020). One control female was omitted from data analyses related to the trauma-paired odor latency because of a time-out response omission. All video behavioral scoring was conducted in BORIS (Friard and Gamba, 2016) by scorers blind to condition and sex of animals.

## ACKNOWLEDGMENTS

We would like to thank Sydney T. Cook & M. Paloma Zacarias for technical assistance. This work was supported by Indiana University – Purdue University Indianapolis. The authors have no conflicts of interest to declare.

## REFERENCES

American Psychiatric Association. 2013. Diagnostic and statistical manual of mental disorders (5th ed.).

Andrews JS. 1996. Possible confounding influence of strain, age and gender on cognitive performance in rats. Brain research. Cognitive Brain Research 3(3-4), 251–267.

April LB, Bruce K, Galizio M. 2013. The Magic Number 70 (plus or minus 20): Variables Determining Performance in the Rodent Odor Span Task. Learning and Motivation 44(3), 143–158.

Ardi Z, Albrecht A, Richter-Levin A, Saha R, Richter-Levin G. 2016. Behavioral profiling as a translational approach in an animal model of posttraumatic stress disorder. Neurobiology of Disease 88:139–47.

Badura-Brack AS, Naim R, Ryan TJ, Levy O, Abend R, Khanna MM, McDermott TJ, Pine DS, Bar-Haim Y. 2015. Effect of Attention Training on Attention Bias Variability and PTSD Symptoms: Randomized Controlled Trials in Israeli and U.S. Combat Veterans. The American Journal of Psychiatry 172(12), 1233–1241.

Balderston NL, Hale E, Hsiung A, Torrisi S, Holroyd T, Carver FW, Coppola R, Ernst M, Grillon C. 2017. Threat of shock increases excitability and connectivity of the intraparietal sulcus. eLife 6, e23608.

Bali A, Jaggi AS. 2015. Electric foot shock stress: a useful tool in neuropsychiatric studies. Reviews in the Neurosciences 26(6):655–77.

Barch DM, Berman MG, Engle R, Jones JH, Jonides J, Macdonald A 3rd, Nee DE, Redick TS, Sponheim SR. 2009. CNTRICS final task selection: working memory. Schizophrenia Bulletin 35(1), 136–152.

Bienvenu T, Dejean C, Jercog D, Aouizerate B, Lemoine M, Herry C. 2021. The advent of fear conditioning as an animal model of post-traumatic stress disorder: Learning from the past to shape the future of PTSD research. Neuron 109(15), 2380–2397.

Block SR, Liberzon I. 2016. Attentional processes in posttraumatic stress disorder and the associated changes in neural functioning. Experimental Neurology 284(Pt B):153–167.

Braun AA, Skelton MR, Vorhees CV, Williams MT. 2011. Comparison of the elevated plus and elevated zero mazes in treated and untreated male Sprague-Dawley rats: effects of anxiolytic and anxiogenic agents. Pharmacology Biochemistry Behavior 97(3):406–15.

Carmassi C, Akiskal HS, Bessonov D, Massimetti G, Calderani E, Stratta P, Rossi A, Dell’Osso L. 2014. Gender differences in DSM-5 versus DSM-IV-TR PTSD prevalence and criteria comparison among 512 survivors to the L’Aquila earthquake. Journal of Affective Disorders 160, 55–61.

Carragher N, Sunderland M, Batterham PJ, Calear AL, Elhai JD, Chapman C, Mills K. 2016. Discriminant validity and gender differences in DSM-5 posttraumatic stress disorder symptoms. Journal of Affective Disorders 190, 56–67.

Cohen H, Kozlovsky N, Alona C, Matar MA, Joseph Z. 2012. Animal model for PTSD: from clinical concept to translational research. Neuropharmacology 62:715–724.

Cohen H, Zohar J, Matar M. 2003. The relevance of differential response to trauma in an animal model of posttraumatic stress disorder. Biological Psychiatry 53(6), 463–473.

Cowan N. 2001. The magical number 4 in short-term memory: a reconsideration of mental storage capacity. The Behavioral and Brain Sciences 24(1), 87–185.

Cowan N. 2008. What are the differences between long-term, short-term, and working memory? Progress in Brain Research 169:323–38.

Cox RC, Olatunji BO. 2017. Linking attentional control and PTSD symptom severity: the role of rumination. Cognitive Behavior Therapy 46(5):421–431.

Davies DA, Greba Q, Selk JC, Catton JK, Baillie LD, Mulligan SJ, Howland JG. 2017. Interactions between medial prefrontal cortex and dorsomedial striatum are necessary for odor span capacity in rats: role of GluN2B-containing NMDA receptors. Learning & Memory 24(10), 524–531.

Davies DA, Molder JJ, Greba Q, Howland JG. 2013. Inactivation of medial prefrontal cortex or acute stress impairs odor span in rats. Learning & Memory 20(12):665–9.

De Brouwer G, Wolmarans W. 2018. Back to basics: A methodological perspective on marble-burying behavior as a screening test for psychiatric illness. Behavioural Processes 157, 590–600.

De Falco E, An L, Sun N, Roebuck AJ, Greba Q, Lapish CC, Howland JG. 2019. The Rat Medial Prefrontal Cortex Exhibits Flexible Neural Activity States during the Performance of an Odor Span Task. eNeuro 6(2), ENEURO.0424-18.2019.

Dudchenko PA. 2004. An overview of the tasks used to test working memory in rodents. Neuroscience and Biobehavioral Reviews 28(7), 699–709.

Dudchenko PA, Talpos J, Young J, Baxter MG. 2013. Animal models of working memory: a review of tasks that might be used in screening drug treatments for the memory impairments found in schizophrenia. Neuroscience and Biobehavioral Reviews 37(9 Pt B), 2111–2124.

Dudchenko PA, Wood ER, Eichenbaum H. 2000. Neurotoxic hippocampal lesions have no effect on odor span and little effect on odor recognition memory but produce significant impairments on spatial span, recognition, and alternation. Journal of Neuroscience 20(8):2964–77.

File SE, Seth P. 2003. A review of 25 years of the social interaction test. European Journal of Pharmacology 463(1-3):35–53.

Fosco WD, Kofler MJ, Groves NB, Chan E, Raiker JS Jr. 2020. Which ‘Working’ Components of Working Memory aren’t Working in Youth with ADHD?. Journal of Abnormal Child Psychology 48(5), 647–660.

Friard O, Gamba M. 2016. BORIS: a free, versatile open-source event-logging software for video/audio coding and live observations. Methods in Ecology and Evolution 7: 1325–1330.

Galizio M, Mason MG, Bruce K. 2020. Successive incrementing non-matching-to-samples in rats: An automated version of the odor span task. Journal of the Experimental Analysis of Behavior 114(2), 248–265.

Gould TD, Dao DT, Kovacsics CE. 2009. The Open Field Test BT. In T.D. Gould (Ed.), Mood and Anxiety Related Phenotypes in Mice: Characterization Using Behavioral Tests (pp.1–20). Humana Press.

Gruene TM, Flick K, Stefano A, Shea SD, Shansky RM. 2015. Sexually divergent expression of active and passive conditioned fear responses in rats. eLife 4:e11352.

Guida A, Gobet F, Tardieu H, Nicolas S. 2012. How chunks, long-term working memory and templates offer a cognitive explanation for neuroimaging data on expertise acquisition: a two-stage framework. Brain and Cognition 79(3), 221–244.

Hahn LA, Balakhonov D, Fongaro E, Nieder A, Rose J. 2021. Working memory capacity of crows and monkeys arises from similar neuronal computations. eLife 10, e72783.

Harvey BH, Naciti C, Brand L, Stein DJ. 2003. Endocrine, cognitive and hippocampal/cortical 5HT 1A/2A receptor changes evoked by a time-dependent sensitisation (TDS) stress model in rats. Brain Research 983(1-2), 97–107.

Himanshu Dharmila, Sarkar D, Nutan. 2020. A Review of Behavioral Tests to Evaluate Different Types of Anxiety and Anti-anxiety Effects. Clinical Psychopharmacology and Neuroscience18(3), 341–351.

Hindley SW, Hobbs A, Paterson IA, Roberts MH. 1985. The effects of methyl beta-carboline-3-carboxylate on social interaction and locomotor activity when microinjected into the nucleus raphé dorsalis of the rat. British Journal of Pharmacology 86(3), 753–761.

Honzel N, Justus T, Swick D. 2014. Posttraumatic stress disorder is associated with limited executive resources in a working memory task. Cognitive, Affective & Behavioral Neuroscience 14(2), 792–804.

Katz RJ, Roth KA, Carroll BJ. 1981. Acute and chronic stress effects on open field activity in the rat: implications for a model of depression. Neuroscience & Biobehavioral Reviews 5(2):247–51.

Kedia S, Chattarji S. 2014. Marble burying as a test of the delayed anxiogenic effects of acute immobilisation stress in mice. Journal of Neuroscience Methods 233, 150–154.

Kessler RC, Berglund P, Demler O, Jin R, Merikangas KR, Walters EE. 2005. Lifetime prevalence and age-of-onset distributions of DSM-IV disorders in the National Comorbidity Survey Replication. Archives of General Psychiatry 62(6):593–602.

Kessler RC, Sonnega A, Bromet E, Hughes M, Nelson CB. 1995. Posttraumatic stress disorder in the National Comorbidity Survey. Archives of General Psychiatry 52(12), 1048–1060.

Khayyer Z, Saberi Azad R, Torkzadeh Arani Z, Jafari Harandi R. 2021. Examining the effect of stress induction on auditory working memory performance for emotional and non-emotional stimuli in female students. Heliyon 7(4), e06876.

Kilpatrick DG, Resnick HS, Milanak ME, Miller MW, Keyes KM, Friedman MJ. 2013. National estimates of exposure to traumatic events and PTSD prevalence using DSM-IV and DSM-5 criteria. Journal of Traumatic Stress 26(5), 537–547.

Knox D, Nault T, Henderson C, Liberzon I. 2012. Glucocorticoid receptors and extinction retention deficits in the single prolonged stress model. Neuroscience 223, 163–173.

Larsen SE, Lotfi S, Bennett KP, Larson CL, Dean-Bernhoft C, Lee HJ. 2019. A pilot randomized trial of a dual n-back emotional working memory training program for veterans with elevated PTSD symptoms. Psychiatry Research 275, 261–268.

Liberzon I, Krstov M, Young EA. 1997. Stress-restress: effects on ACTH and fast feedback. Psychoneuroendocrinology 22(6), 443–453.

Martis LS, Krog S, Tran TP, Bouzinova E, Christiansen SL, Møller A, Holmes MC, Wiborg O. 2018. The effect of rat strain and stress exposure on performance in touchscreen tasks. Physiology & Behavior 184, 83–90.

McDermott TJ, Badura-Brack AS, Becker KM, Ryan TJ, Bar-Haim Y, Pine DS, Khanna MM, Heinrichs-Graham E, Wilson TW. 2016. Attention training improves aberrant neural dynamics during working memory processing in veterans with PTSD. Cognitive, Affective, & Behavioral Neuroscience 16(6):1140–1149.

Meyerson BJ, Augustsson H, Berg M, Roman E. 2006. The Concentric Square Field: a multivariate test arena for analysis of explorative strategies. Behavioural Brain Research 168(1), 100–113.

Mikics E, Baranyi J, Haller J. 2008. Rats exposed to traumatic stress bury unfamiliar objects--a novel measure of hyper-vigilance in PTSD models? Physiology & Behavior 94(3):341–8.

Miller GA. 1956. The magical number seven plus or minus two: some limits on our capacity for processing information. Psychological Review 63(2), 81–97.

Moran TP. 2016. Anxiety and working memory capacity: A meta-analysis and narrative review. Psychological Bulletin 142(8):831–864.

Murphy S, Elklit A, Chen YY, Ghazali SR, Shevlin M. 2019. Sex differences in PTSD symptoms: A differential item functioning approach. Psychological Trauma: Theory, Research, Practice and Policy 11(3), 319–327.

Nejati V, Salehinejad MA, Sabayee A. 2018. Impaired working memory updating affects memory for emotional and non-emotional materials the same way: evidence from post-traumatic stress disorder (PTSD). Cognitive Processing 19(1), 53–62.

Peters A, McEwen BS, Friston K. 2017. Uncertainty and stress: Why it causes diseases and how it is mastered by the brain. Progress in Neurobiology 156, 164–188.

R Core Team. 2020. R: A language and environment for statistical computing. R Foundation for Statistical Computing, Vienna, Austria. URL https://www.R-project.org/.

Richter-Levin G, Stork O, Schmidt MV. 2019. Animal models of PTSD: a challenge to be met. Molecular Psychiatry 24(8):1135–1156.

Sánchez-Andrade G, James BM, Kendrick KM. 2005. Neural encoding of olfactory recognition memory. The Journal of Reproduction and Development 51(5), 547–558.

Saunders N, Downham R, Turman B, Kropotov J, Clark R, Yumash R, Szatmary A. 2015. Working memory training with tDCS improves behavioral and neurophysiological symptoms in pilot group with post-traumatic stress disorder (PTSD) and with poor working memory. Neurocase 21(3), 271–278.

Schoofs D, Pabst S, Brand M, Wolf OT. 2013. Working memory is differentially affected by stress in men and women. Behavioural Brain Research 241, 144–153.

Scott JC, Matt GE, Wrocklage KM, Crnich C, Jordan J, Southwick SM, Krystal JH, Schweinsburg BC. 2015. A quantitative meta-analysis of neurocognitive functioning in posttraumatic stress disorder. Psychological Bulletin 141(1):105–140.

Shepherd JK, Grewal SS, Fletcher A, Bill DJ, Dourish CT. 1994. Behavioural and pharmacological characterisation of the elevated “zero-maze” as an animal model of anxiety. Psychopharmacology 116(1):56–64.

Shields GS, Sazma MA, Yonelinas AP. 2016. The effects of acute stress on core executive functions: A meta-analysis and comparison with cortisol. Neuroscience & Biobehavioral Reviews 68:651–668.

Spataro P, Mulligan NW, Saraulli D, Rossi-Arnaud C. 2022. The attentional boost effect facilitates the encoding of contextual details: New evidence with verbal materials and a modified recognition task. Attention, Perception & Psychophysics 10.3758/s13414-022-02509-z. Advance online publication.

Tucker LB, McCabe JT. 2017. Behavior of Male and Female C57BL/6J Mice Is More Consistent with Repeated Trials in the Elevated Zero Maze than in the Elevated Plus Maze. Frontiers in Behavioral Neuroscience 11:13.

Turchi J, Sarter M. 2000. Cortical cholinergic inputs mediate processing capacity: effects of 192 IgG-saporin-induced lesions on olfactory span performance. The European journal of Neuroscience 12(12), 4505–4514.

Van Dijken HH, Van der Heyden JA, Mos J, Tilders FJ. 1992. Inescapable footshocks induce progressive and long-lasting behavioural changes in male rats. Physiology & Behavior 51(4):787–94.

Verbitsky A, Dopfel D, Zhang N. 2020. Rodent models of post-traumatic stress disorder: behavioral assessment. Translational Psychiatry 10(1), 132.

Voyer D, Saint Aubin J, Altman K, Gallant G. 2021. Sex differences in verbal working memory: A systematic review and meta-analysis. Psychological Bulletin 147(4), 352–398.

Wellman LL, Fitzpatrick ME, Machida M, Sanford LD. 2014. The basolateral amygdala determines the effects of fear memory on sleep in an animal model of PTSD. Experimental Brain Research 232(5):1555–65.

Woodward SH, Jamison AL, Gala S, Holmes TH. 2017. Canine companionship is associated with modification of attentional bias in posttraumatic stress disorder. PloS One 12(10), e0179912.

Yehuda R, Antelman SM. 1993. Criteria for rationally evaluating animal models of posttraumatic stress disorder. Biological Psychiatry 33(7):479–86.

